# Analysis of sex-specific lipid metabolism in *P. falciparum* gametocytes points to importance of sphingomyelin for gametocytogenesis

**DOI:** 10.1101/2021.11.02.467029

**Authors:** Melanie C. Ridgway, Daniela Cihalova, Simon H.J. Brown, Phuong Tran, Todd W. Mitchell, Alexander G. Maier

## Abstract

Male and female *Plasmodium falciparum* gametocytes are the parasite lifecycle stage responsible for transmission of malaria from the human host to mosquito vector. Not only are gametocytes able to survive in radically different host environments, but they are also precursors for male and female gametes that reproduce sexually soon after ingestion by the mosquito. Here we investigate the sex-specific lipid metabolism of gametocytes within their host red blood cell and poised for ingestion by the mosquito vector and subsequent sexual reproduction.

Comparison of the male and female lipidome identifies cholesteryl esters and dihydrosphingomyelin enrichment in female gametocytes. Chemical inhibition of each of these lipid types in mature gametocytes suggests dihydrosphingomyelin synthesis but not cholesteryl ester synthesis is important for sex-specific gametocyte viability. Genetic disruption of each of the two sphingomyelin synthase gene points towards sphingomyelin synthesis contributing to gametocytogenesis.

This study shows that gametocytes are not only distinct from asexual stages, but that the lipid composition is also vastly different between male and female gametocytes, reflecting the different cellular roles these stages play. Together our results highlight the sex-specific nature of gametocyte lipid metabolism that has the potential to be targeted to block malaria transmission.

## Introduction

Transmission of *Plasmodium falciparum* from the human to mosquito host depends on male and female gametocytes. The development of transmissible gametocytes (gametocytogenesis) is by far the longest developmental sexual stage in *P. falciparum*. After 10-12 days of maturation in the human red blood cell (RBC), ingestion by a mosquito rapidly activates the gametocytes to form mature gametes capable of sexual reproduction (Bennink et al. 2016). Within 15 minutes of the blood meal, gametocytes egresses from their host RBC. While male gametocytes multiply into eight flagellated microgametes, female gametocytes form an immobile spherical macrogamete, poised for fertilisation. The zygote develops in the extracellular environment until invasion of the mosquito midgut wall 19-36 hours post blood meal (Aikawa et al. 1984; Sinden et al. 1985; Vlachou et al. 2004).

Such a rapid sex-specific metamorphosis is necessarily preceded by sex-specific preparations in the human host during the relatively slow gametocytogenesis. Surprisingly though, morphological gametocyte sex dimorphism is relatively subtle and mostly limited to the ultrastructural level. Recently the tagging of sex-specific molecular markers has revealed sex-specific gene and protein expression profiles (Lasonder et al. 2016; Miao et al. 2017). These studies identified differences in the expression of mRNA and protein involved in lipid metabolic pathways including phosphatidylcholine biosynthesis, fatty acid biosynthesis and ether lipid metabolism (Lasonder et al. 2016). Whether this results in differences in the lipid composition between male and female gametocytes remains unknown.

Lipids fulfil a vast range of functions in the cell. They not only provide building blocks in the form of fatty acids for the parasite’s metabolism, but they also form the basis for membranes (mainly phospholipids and cholesterol), functionalise subdomains within membranes (e.g. sphingolipids), serve as energy storage (e.g. neutral lipids like cholesterylesters and triacylglycerol) and act as signalling molecules (e.g. diacylglycerol and ceramide). The nature of lipids in male and female gametocytes may reveal diverging cellular functions that precede sex-specific development in the mosquito.

Based on their resilience against many drugs that are active against asexual forms, gametocytes were generally thought to be relatively metabolically inactive (Canning & Sinden 1975; Sinden et al. 1978). However, the lipidome of combined male and female gametocytes changes significantly between young and mature gametocytes (Gulati et al. 2015; Tran et al. 2016), suggesting that lipid metabolism is active during gametocytogenesis. In particular, Gulati et al. (2015) identified intermediate products of sphingolipid metabolism in mature gametocytes, revealing that *de novo* sphingomyelin synthesis occurs during gametocytogenesis.

Sphingomyelin is synthesised *de novo* from serine and palmitoyl-CoA in five consecutive reactions. The final step of *de novo* sphingomyelin synthesis is mediated by sphingomyelin synthase (SMS). *P. falciparum* encodes two SMS (Gardner et al. 2002), one of which is expressed throughout blood stages (SMS1, PF3D7_0625000), the other of which is gametocyte specific and female enriched (SMS2, PF3D7_0625100) (López-Barragán et al. 2011; Lasonder et al. 2016). In addition to generating sphingomyelin, SMS also regulates the levels of secondary messenger molecules such as ceramide and diacylglycerol (Fyrst & Saba 2010). The functional significance of SMS1 and SMS2 in *P. falciparum* has yet to be investigated.

Here we explore the sex-specific lipidome of mature *P. falciparum* gametocytes within their host RBC to determine the respective contribution of male and female gametocytes to the lipid profile of the infected RBC (iRBC). The functional significance of *de novo* synthesis of key sex-specific lipids is then investigated by pharmacological inhibition and reverse genetics.

## Materials and Methods

### Culturing techniques

Asexual 3D7 strain *P. falciparum* parasites were maintained in complete culture medium (RPMI 1640-Hepes with Glutamax media supplemented with 10 mM D-glucose, 480 *μ*M hypoxanthine, 20 *μ*g. mL gentamicin, 0.375% (w/v) Albumax II and 2.5% v/v heat-inactivated human serum) as previously described (Maier & Rug 2013). Cultures were synchronised by sorbitol treatment as described previously (Lambros & Vanderberg 1979) and isolated from their host cells by saponin treatment (Dourmashkin et al. 1962).

Parasites were induced to form gametocytes as described by Fivelman et al. (2007) with modifications to reduce asexual parasite proliferation (Ridgway et al. 2020). Male and female gametocytes were collected by fluorescence activated cell sorting as described by Ridgway et al. (2021). Briefly, magnet enriched 3D7 gametocytes with a female-specific GFP tag (gABCG2-GFP, (Tran et al. 2014)) were stained with Hoechst 33342 and collected by fluorescence activated cell sorting on day 9 post commitment (mostly at stage IV). Hoechst staining identified gametocyte-infected from uninfected RBC, while the female-specific GFP signal distinguished male and female gametocytes.

Gametocytemia was measured every second day from day 2 post commitment by flow cytometry of an aliquot of culture stained with 50 μg/mL Hoechst 33342 in PBS (10.6 mM KH_2_PO4, 1.55 M NaCl, 29.7 mM Na_2_HPO4, pH 7.4) for 15 min at 37°C, rinsed twice in PBS with 2,000 ×*g*, 1 min spins and resuspended in PBS. Samples were analysed on a LSR II Flow Cytometer (BD Biosciences) detecting Hoechst 33342 in the Pacific Blue channel (350 nm excitation/461 nm emission). Parasitemia was calculated as the proportion of Hoechst 33342 positive cells in 200,000 whole, single cells as gated in FACS Diva software. Statistical significance and graphing of results was performed in GraphPad Prism 7.

### Sex-specific gametocyte viability assay

Magnet enriched gametocytes were exposed to 10 μM of each test compound, 200 μM artemisinin (0% viable control) or 0.1% (v/v) DMSO (100% viable control) for 96 h from day 2 to 6 post commitment in complete culture medium with 50 mM N-acetyl D-glucosamine in a 96 well plate at 37°C in hypoxic conditions (1% O_2_, 5% CO_2_, 94% N_2_).

Parasites were then stained in 50 μg/mL Hoechst 33342 and 500 nM MitoTracker Deep Red in complete culture medium supplemented with 50 mM N-acetyl D-glucosamine for 30 min at 37 °C in hypoxic conditions (1% O_2_, 5% CO_2_, 94% N_2_). Stained cells were rinsed twice in PBS with 1,000 ×*g*, 5 min spins and resuspended in PBS. Single colour controls for flow cytometry consisted of i) asexual 3D7 WT culture stained with 500 nM MitoTracker Deep Red or 50 μg/mL Hoechst 33342 for 30 min in complete culture medium, rinsed twice in PBS with 1,000 ×*g*, 1 min spins and resuspended in PBS, ii) unstained 3D7 gABCG2-GFP gametocytes suspended in PBS and iii) unstained asexual 3D7 WT parasites in PBS (for the unstained control). Samples were analysed on a LSR II Flow Cytometer (BD Biosciences) detecting Hoechst 33342 in the Pacific Blue channel (405 nm excitation/461 nm emission), MitoTracker Deep Red in the APC-Cy7 channel (644 nm excitation/665 nm emission) and GFP in the FITC channel (488 nm excitation/509 nm emission). A total of 500,000 events were recorded in each single colour control. In each sample data was recorded until 500,000 female gametocytes were counted.

After applying the gating strategy to isolate male and female gametocytes (Ridgway et al. 2020) the mean fluorescence intensity of MitoTracker Deep Red detected on the APC-Cy7 channel was recorded for each of the gametocyte populations. Data was analysed in FlowJo and statistical significance was determined by ANOVA (analysis of variance) in GraphPad Prism. Cell viability is expressed as a percentage of the mean fluorescence intensity of MitoTracker Deep Red in gametocytes treated with 0.1% DMSO (100% viable) and 200 μM artemisinin treated gametocytes (0% viable).

### Generation of transgenic *P. falciparum* cell lines

Annotated sequence of *P. falciparum* 3D7 reference strain was sourced from the *Plasmodium* Genomics Resource PlasmoDB (www.plasmodb.org). To disrupt SMS1 (PF3D7_0625000) by double recombination, a 5’ fragment and a 3’ fragment of SMS1 locus were amplified using primers al398/al399 and al400/al401 (Supplementary Table 1) and cloned into a pCC-1 vector containing a human dihydrofolate reductase (hDHFR) and 5-fluorocytosine drug selection cassette (Maier et al. 2008) at SacII/SpeI and EcoRI/AvrII sites to generate pCC-1/SMS1 construct (Supplementary Figure 1). SMS2 (PF3D7_0625100) was also disrupted by double recombination by amplifying a 5’ fragment with al406/al407 primers and a 3’ fragment with al408/409 primers (Supplementary Table 1), which were cloned into a pCC-1 vector at SacII/SpeI and EcoRI/AvrII sites to generate pCC-1/SMS2 construct (Supplementary Figure 2). Given that SMS1 and SMS2 are adjacent genes, the pCC-1/SMS1 and 2 construct aiming to disrupt both SMS genes was generated by cloning the 5’ fragment of SMS1 (amplified with al398/al399 primers) and the 3’ fragment of SMS2 (amplified with al408/al409 primers) at SacII/SpeI and EcoRI/AvrII sites. Inserted DNA regions were confirmed by analytical restriction enzyme digest and sequenced to verify constructs. Plasmids were purified using Invitrogen Purelink Maxiprep kit prior to transfection.

*P. falciparum* 3D7 reference strain parasites were transfected as described previously (Rug & Maier 2013). Briefly, 400 μL of 100 μg of plasmid DNA in Cytomix (120 mM KCl, 0.15 mM CaCl_2_, 10 mM K_2_HPO_4_/KH_2_PO_4_, 25 mM HEPES, 2 mM ethylene glycol-bis(β-aminoethyl ether)-tetraacetic acid, 5 mM MgCl_2_, pH 7.6) was electroporated into a sorbitol synchronised ring stage culture at 5% parasitemia by 310 V, 950 μF pulse delivered in a 0.2 cm electrode gap cuvette placed in a BioRad Genepulser II with ∞ capacitor. Transformed cells were immediately resuspended in complete culture medium with 3% haematocrit uninfected RBC and incubated at 37°C in hypoxic conditions (1% O_2_, 5% CO_2_, 94% N_2_). Culture medium was supplemented with 2 nM WR99210 from 4 h post transfection and was replaced daily for 5 days post transfection, then three times a week until parasites were observed by Giemsa-stained thin smears of the culture. Parasites having integrated the plasmid were enriched by three 21-day off/on 2 nM WR99210 cycles then negative selection was applied by adding 231 nM 5-fluorocytosine. The transgenic parasites were cloned by limited dilution (Rug & Maier 2013).

### DNA extraction

Genomic DNA was extracted from saponin-isolated trophozoite stage parasites using DNeasy Blood and Tissue kit (Qiagen) as per the manufacturer’s instructions. For Southern blot analysis, genomic DNA was eluted twice in 200 μL elution buffer each then concentrated by adding 40 μL 3 M sodium acetate and 880 μL 100% ethanol at 4°C overnight. Samples were centrifuged at 17,000 ×*g* for 30 min at 4°C then DNA was washed twice in 70% ethanol with 17,000 ×*g*, 20 min spins. DNA was air dried then re-suspended in TE buffer (10 mM Tris base, 1 mM EDTA, pH 8.0). DNA concentration and purity was measured by NanoDrop spectrophotometer (Thermofisher).

### Quantitative PCR

The abundance of WT and knock out (KO) genotypes in co-cultures was monitored by quantitative PCR (qPCR) of regions of DNA specific to each genome. SMS1 KO and SMS2 KO parasites were each quantified by primers specific to the sequence of the inserted hDHFR drug resistance cassette. WT parasites were distinguished from SMS1 KO parasites by primers specific to the region of SMS1 disrupted by homologous recombination in the SMS1 KO parasites. WT parasites in the SMS2 KO/WT co-cultures were similarly quantified by primers specific to the disrupted section of SMS2 in the SMS2 KO parasites. The primers for this experiment are described in Supplementary Table 2.

The qPCR were performed using a Light Cycler 480 SYBR Green I Master mix (Roche) as per the manufacturer’s instructions for 10 μL reactions in a 384 well plate. Thermocycling was performed with a 10 min, 95°C pre-incubation; 45 cycles of 15 s denaturation at 95°C, 15 s annealing at 52°C and 20 s elongation at 72°C; followed by a melt curve established by denaturing at 95°C for 30 s, annealing at 60°C for 30 s and then slowly denaturing by increasing the temperature to 95°C at 0.11°C/s. Melt curves were observed using Light Cycler 480 software. The exported text file was converted using Light Cycler 480 converter (Roche) and Cq and PCR efficiencies were determined using the LinReg program (Ruijter et al. 2009). Relative quantification of transcripts was expressed as described previously (Pfaffl 2001).

For the quantification of WT and KO parasite genotypes in co-cultures:

ratio = (E_target_^ΔCPtarget(clonal-competition)^)/(E_ref_^ΔCPref(clonal-competition)^)
with: E_target_: primer efficiency of the target gene E_ref_: primer efficiency of the reference gene ΔCP_target_ (clonal-competition): difference in crossing points 225 of target gene amplification in a clonal population and the fitness competition co-culture ΔCP_ref_ (clonal-competition): difference in crossing points of reference gene amplification in a clonal population and the fitness competition co-culture

### Southern Blot

Disruption of a gene locus by homologous recombination was confirmed by diagnostic digest with restriction enzymes followed by Southern blot probing for both the 5’ and 3’ homologous regions (Rug & Maier 2013). DNA was digested with either AflII and PacI (for SMS1 KO screening) or Xmn I and Hind III-HF (for SMS2 KO screening) as per the manufacturer’s instructions then run on a 0.8% agarose gel at 90 V for 15 min followed by a run at 20 V for 18 h. The gel was then rinsed as follows: 15 min in depurination solution (0.25 M HCl); 3 min in milliQ water; 2*×* 15 min in denaturation solution (0.5 M NaOH, 1.5 M NaCl); 3 min in milliQ water; 2*×* 15 min in neutralisation solution (0.5 M Tris base, 1.5 M NaCl, pH 7.5); 3 min in milliQ water then 5 min in 20× SCC (3 M NaCl, 0.3 M citric acid trisodium dihydrate, pH 7.0). DNA was transferred from the gel to the membrane overnight in a capillary transfer setup.

Alkali-labile digoxigenin labelled deoxyuridine triphosphate (DIG-dUTP) was incorporated in Southern blot probes by PCR using PCR DIG Probe Synthesis kit (Roche) with the following specifications (refer to sequences in Supplementary table 3). Each probe was amplified from the corresponding plasmid using either OneTaq (NEB) or Taq polymerase (Roche). SMS1 probes were purified by gel extraction (Qiagen kit). All probes were synthesised in the presence of 12 μM DIG-dUTPs except the SMS2 3’ probe synthesised from 6 μM DIG-dUTPs. Thermocycling conditions consisted of initial denaturation for 30 s at 94°C; 30 cycles of 30 s at 94°C, 1 min at 45°C (SMS2 3’) or 50°C (SMS2 5’) or 52°C (SMS1 5’) or 55°C (SMS1 3’), 1 min at 68°C; final elongation for 7 min at 68°C.

Following transfer, the membrane was rinsed in 2*×* SCC for 5 min then DNA was crosslinked to the membrane in a UV cross linker set to deliver 70 mJ/cm^2^. The membrane was incubated with gentle rocking in DIG Easy Hyb solution (Roche Cat. No. 11 796 895 001) at the respective probe hybridisation temperature (T_hyb_) for 30 min. The probe was denatured in 50 μL water at 95°C for 5 min, cooled quickly on ice then added to DIG Easy Hyb solution at 0.15% (v/v) to prepare the hybridisation solution. The membrane was incubated overnight in hybridisation solution at T_hyb_ with gentle rocking.

The membrane was prepared for detection by rinsing in 2× SCC for 5 min; equilibrating in washing buffer for 1 min (0.1 M maleic acid, 0.15 M NaCl, pH 7.5 with 0.3% (v/v) Tween-20); incubating in 1% blocking solution (0.1 M maleic acid, 0.15 M NaCl, pH 7.5 with 1% (w/v) skim milk powder) for 30 min; incubating in 1% blocking solution with 1:10,000 α-Digoxigenin-AP Fab fragments (Roche) for 30 min; washing membrane in washing buffer for 2× 15 min then equilibrating membrane in detection buffer (100 mM Tris-hydrochloride, 100 mM NaCl, pH 9.5) for 2 min. The membrane was incubated in the dark with CSPD (Roche) diluted 1:100 in detection buffer for 5 min at room temperature then 10 min at 37°C. The membrane was exposed to Fuji Super RX-N Medical X-Ray film in an Amersham pharmacia biotech hypercassette and developed film in an AGFA CP1000 photo developer.

### Lipidomic analysis

Uninfected human RBC from six donors were pooled in three independent biological replicates each from two donors and incubated at 4% haematocrit in complete culture medium for at least 48 h at 37°C in hypoxic conditions (1% O_2_, 5% CO_2_, 94% N_2_). Cells were pelleted at 524 ×*g* over 5 min and counted on an Improved Neubauer haemocytometer (Hirschmann). Aliquots of 10^7^ cells were resuspended in 300 μL of methanol in 2 mL tough tubes (Geneworks) and stored at −80°C. For each biological replicate parasites of different stages where grown in the same donor batch of RBC to reduce host specific influences.

For collection of asexual parasite infected RBC, three independent asexual *P. falciparum* cultures were sorbitol synchronised then magnet purified to more than 98% iRBC, as determined by Giemsa-stained thin smear, and counted on an Improved Neubauer haemocytometer (Hirschmann). Aliquots of 10^7^ iRBC at 26-42 h post invasion were resuspended in 300 μL of methanol in 2 mL tough tubes (Geneworks) and stored at −80°C.

Male and female gametocytes were sorted live by FACS as described above, pelleted at 754 ×*g* for 10 min and resuspended in 300 μL of methanol per 10^7^ cells (as counted during FACS) in 2 mL tough tubes (Geneworks). Each biological replicate of 10^7^ cells is pooled from 2-5 independent gametocyte cultures.

Lipids were extracted as described previously (Matyash et al. 2008) with modifications. Internal standards (Tran et al. 2016) in 50 μL of methanol with 0.01% butylated hydroxytoluene were aliquoted among tubes. Internal standards and 1.4 mm beads (Geneworks) were added to tough tubes containing samples in 300 μL methanol and samples were homogenized with a bead homogenizer (FastPrep-24, MP Biomedical) at 6 m/s for 40 s and transferred to new tubes. Beads were rinsed in 100 μL of methanol, which was added to the sample in new tubes. Samples were vortexed with 1 mL of methyl-tert butyl ether at 4°C for 1 h. Phase separation was induced by adding 300 μL of 150 mM ammonium acetate (liquid chromatography–MS grade, Fluka), vortexing for 5 min and centrifuging at 2,000 ×*g* for 5 min. The upper organic layer (representing around 800 μL) was transferred to a 2 mL glass vial and stored at −20°C. Prior to MS analysis, each sample was diluted into methanol:chloroform (2:1 v/v) with 5 mM ammonium acetate.

An aliquot of each extract was hydrolyzed to remove acyl-linked lipids and re-extracted to improve mass spectrometric analysis of sphingolipids. Two hundred μL of extract was added to 60 μL of methanol containing 0.01% butylated hydroxytoluene, 22 μL of 10 M NaOH was added (final concentration 0.7 M), and vortexed at 800 rounds per minute on an Eppendorf Mixmate for 2 h at room temperature. Sixty μL of 150 mM aqueous ammonium acetate was added to induce phase separation. Tubes were vortexed and spun at 20,000 ×*g* for 5 min to complete phase separation. The upper organic layer was removed to a new 2 mL glass vial, and diluted into methanol:chloroform (2:1 v/v) containing 5 mM ammonium acetate prior to mass spectrometric analysis.

Mass spectra were obtained with a chip-based nanoESI source (TriVersa Nanomate, Advion) and a hybrid linear ion-trap-triple quadrupole MS (QTRAP 5500, ABSCIEX). Ten μL of each sample extract in a sealed Eppendorf Twin-Tec 96 well plate was aspirated and delivered to the MS through a nanoESI chip. Positive ion and negative ion acquisition was obtained as described previously (Tran et al. 2016).

Data smoothing, lipid identification, removal of isotope contribution from lower mass species and correction for isotope distribution was performed in LipidView (ABSCIEX) software version 1.2. A signal to noise ratio threshold of 20 was applied for inclusion of ionized lipids. Extraction and solvent blanks were analyzed in each data acquisition batch to exclude chemical or solvent impurities. Lipids were quantified in LipidView by comparing peak area of each lipid to its class-specific internal standard after isotope correction. Where odd-chain fatty acid phospholipids or ether-linked phospholipids could not be distinguished, phospholipids were assumed to be ether linked. A correction factor of 3.45 was applied to all ether-PE species to account for the ~29% difference in efficiency in the neutral loss of the 141 Da fragment from plasmenyl and diacyl PE respectively (Mitchell et al. 2007; Abbott et al. 2013). Lipid species are annotated as per Liebisch et al. (2013) shorthand, except for DAG and TAG.

LipidView data was exported to Excel then imported to Markerview v1.2.1.1 (Applied Biosystems, MDS Sciex) with lipid annotation and quantification for statistical analysis of all individual lipid species. Pareto scaling without weighting was applied for unsupervised mode principal component analysis. Multiple t-tests for volcano plots were also performed in MarkerView (MDS Sciex). Grouping into lipid classes and calculation of lipid proportions were performed in Microsoft Excel. GraphPad Prism 7 was used to prepare graphs and perform ANOVA with multiple comparisons on lipid class abundance.

## Results

### The lipid composition of infected RBC is characteristic of the parasite’s sex and lifecycle stage

The lipidomic analysis was performed on uninfected RBC and RBC infected with male gametocytes, female gametocytes or mature asexual blood stage parasites. Over 230 lipid species were identified in the samples (for a complete list refer to Supplementary Table 4). We performed a principle component analysis to determine the relative contribution of each of these lipid species to the variation between samples (Figure 1). The principal component analysis plot (Figure 1A) groups together samples of similar lipid composition and the corresponding loading plot (Figure 1B) illustrates which lipid species distinguish sample groups.

**Figure 1:**
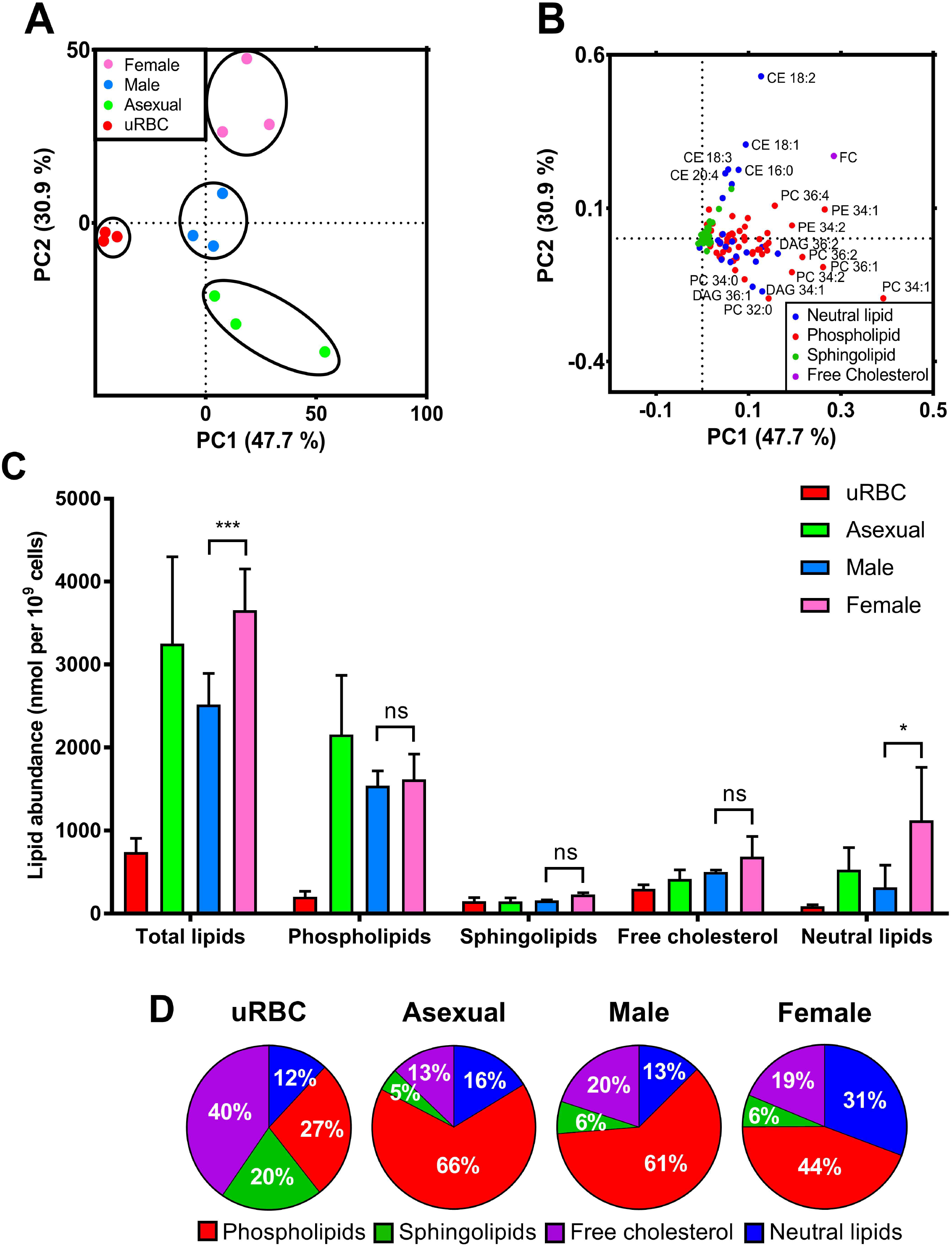
A-B: Principal component analysis (A) and loading plot (B) of the lipidome of uninfected red blood cells (uRBC-red) and red blood cells infected with asexual stage parasites (Asexual-green), female gametocytes (Female - pink) and male gametocytes (Male - blue). Black circles in A indicate sample groups. The individual lipid species are colour-coded to indicate the group they belong to (Neutral lipids – blue; Phospholipids – red, Sphingolipids – green, Free Cholesterol – indigo). C-D: Mean lipid abundance with standard deviation (C) and proportions (D) of lipid classes in uninfected red blood cells (uRBC) and red blood cells infected with asexual stage parasites (Asexual), male gametocytes (Male) and female gametocytes (Female). Results averaged from three independent biological replicates of 10^7^ cells each. Significance calculated by ANOVA. ***: p<0.001, *: p<0.1, ns: not significant, p> 0.1. PC1 and 2: first and second principal component; CE: cholesteryl ester; FC: free cholesterol; PC: phosphatidylcholine; PE: phosphatidylethanolamine; DAG: diacylglycerol.

In considering all measured lipids by principal component analysis, the biggest difference in lipid composition distinguishes infected from uninfected RBC regardless of parasite lifecycle stage (principal component (PC) 1, Figure 1A). A third of the lipid variation between samples nevertheless separates asexual parasite-infected RBC, male gametocyte-infected RBC and female gametocyte-infected RBC (principal component (PC) 2, Figure 1A). Parasite stages mainly cluster based on cholesteryl ester (CE) species, free cholesterol (FC), diacylglycerol (DAG) and phosphatidylcholine (PC) (Figure 1B).

### Female gametocyte infected RBC contain more neutral lipids than other stages

Upon infection, total lipid abundance in infected RBC increases 3-5 fold and in sexual stages, female gametocyte-infected RBC accumulate significantly more lipids than male gametocyte-infected RBC (Figure 1C). However, this increase is not reflected across all lipids: of the four main lipid categories (phospholipids, sphingolipids, free cholesterol and neutral lipids), only neutral lipids are significantly more abundant in female gametocyte-infected RBC compared to male gametocyte-infected RBC (Figure 1C). This is also reflected in the relative proportions of lipid groups in each stage: neutral lipids account for 31% of lipids in female gametocyte-infected RBC compared to only 13% in male gametocyte-infected RBC (Figure 1D). Sphingolipid and free cholesterol contents are similar in abundance but decreases in relative terms in RBCs upon infection or in sexual stages. Phospholipids on the other hand are more abundant and represent a larger proportion to the relative lipid contribution of infected RBCs.

In general, neutral lipids are a cellular means of storing energy and contribute to intracellular signalling pathways. The nature of neutral lipids in asexual parasite-infected RBC contrasts to that of gametocyte-infected RBC (Figure 2). Of the three neutral lipids groups (cholesteryl ester (CE), diacylglycerol (DAG) and triacylglycerol (TAG)), DAG and TAG are the predominant neutral lipids in asexual parasite-infected RBC, whereas CE dominates the neutral lipid profile of gametocyte-infected RBC and uninfected RBC (Figure 2A and B). The proportions of neutral lipid categories are highly dynamic upon infection with asexual or sexual stage *P. falciparum*. In addition, accumulation of neutral lipids is sex-specific: female gametocyte-infected RBC accumulate significantly more CE than male gametocyte-infected RBC (Figure 2A). These trends are also observed at the individual lipid species level (Figure 2C and D). CE species characterise female gametocyte-infected RBC while asexual parasite-infected RBC are distinguished by DAG and TAG species (Figure 2C and D).

**Figure 2:**
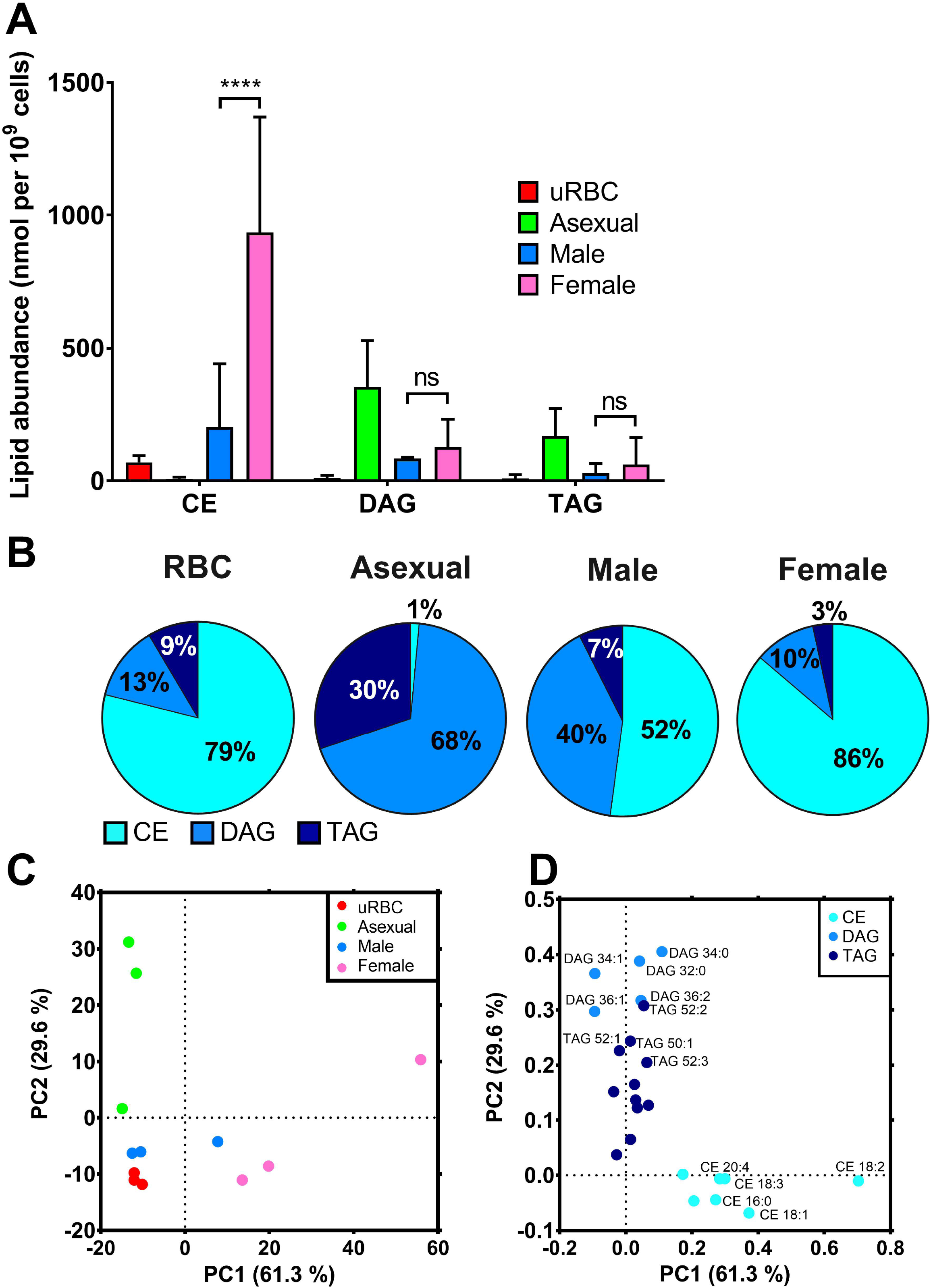
A-B: Mean neutral lipid abundance with standard deviation (A) and mean proportions (B) of uninfected red blood cells (uRBC) and red blood cells infected with asexual stage parasites (Asexual), male gametocytes (Male) or female gametocytes (Female). Results averaged from three independent biological replicates of 10^7^ cells. Significance calculated by ANOVA. ****: p<0.0001, ns: not significant, p> 0.1. CE: cholesteryl ester; DAG: diacylglycerol; TAG: triacylglycerol. C-D: Principal component analysis (C) and loading plot (D) of neutral lipid composition of uninfected red blood cells (uRBC) and red blood cells infected with asexual stage parasites (Asexual), male gametocytes (Male) or female gametocytes (Female). PC1 and 2: first and second principal component; CE: cholesteryl ester; DAG: diacylglycerol; TAG: triacylglycerol.

### Phospholipids of gametocyte-infected RBC differ from those of asexual stage parasite infected RBC but are not sex-specific

Phospholipids are the main component of membranes and amongst others are required for organelle biogenesis. Hence it is not surprising that most lipids in iRBC belong to this group (Figure 1A). Although male gametocyte-infected RBC have a higher proportion of phospholipids (61% phospholipids in males compared to 44% in females, Figure 1D), phospholipid abundance is similar in male and female gametocyte-infected RBC (Figure 1C). There were no significant differences in major phospholipid groups phosphatidyl choline (PC), phosphatidyl ethanolamine (PE), phosphatidyl serine (PS) or phosphatidyl glycerol (PG) between sexes (Figure 3A). PG is only detected in iRBC. Male gametocyte-infected RBC contain slightly more PC but slightly less PE than female gametocyte-infected RBC, both in terms of abundance and proportion of phospholipids (Figure 3A and B).

**Figure 3:**
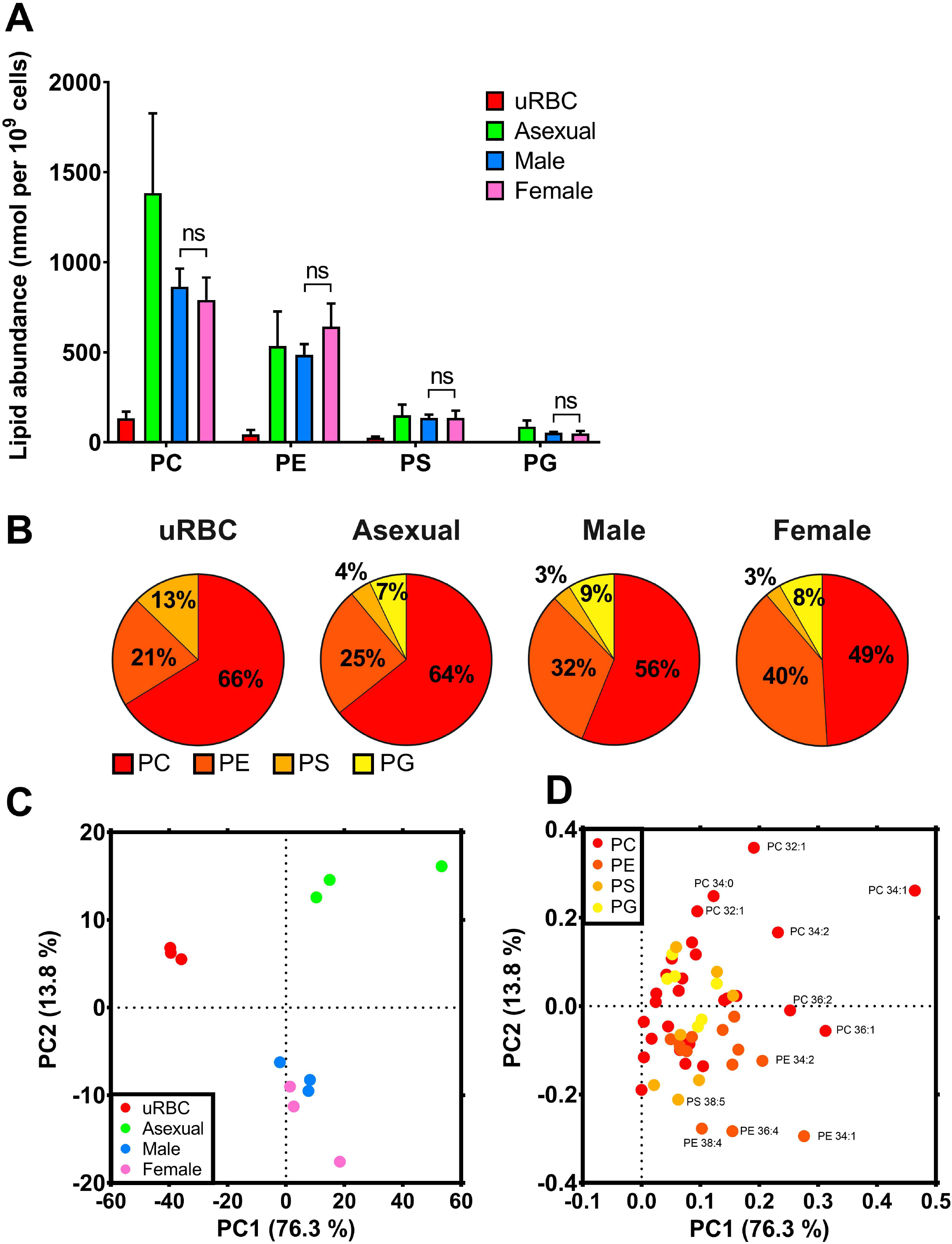
A-B: Mean phospholipid composition with standard deviation (A) and mean proportions (B) of uninfected red blood cells (uRBC) and red blood cells infected with asexual stage parasites (Asexual), male gametocytes (Male) or female gametocytes (Female). Results averaged from three independent biological replicates of 10^7^ cells. Significance calculated by two-way ANOVA. ns: not significant, p> 0.1. C-D: Principal component analysis (C) and loading plot (D) of phospholipid composition of uninfected red blood cells (uRBC) and red blood cells infected with asexual stage parasites (Asexual), male gametocytes (Male) or female gametocytes (Female). PC1 and 2: first and second principal component; PC: phosphatidylcholine; PE: phosphatidylethanolamine; PS: phosphatidylserine; PG phosphatidylglycerol.

Principal component analysis based only on phospholipids does not distinguish between male and female gametocyte-infected RBC (Figure 3C). However, gametocyte-infected RBC cluster apart from uninfected RBC and asexual parasite-infected RBC mostly due to PE species (Figure 3C and D). Overall, male and female gametocyte-infected RBC phospholipids are similar to each other, but distinct from host cell and asexual parasite-infected RBC phospholipids.

### The sphingolipid dihydrosphingomyelin is a key characteristic of female gametocytes

Sphingolipids are also components of cell membranes and form detergent-resistant lipid domains that are platforms for intracellular signalling. Like phospholipids, overall sphingolipid abundance is not significantly different between male and female gametocyte-infected RBC (Figure 1C). However, breaking down sphingolipids into its three main groups (sphingomyelin (SM), dihydrosphingomyelin (DHSM) and ceramide) reveals gametocyte and sex-specific sphingolipid groups (Figure 4). Unlike the abundance of the major sphingolipid SM that is similar in all samples (Figure 4A), gametocyte-infected RBC (especially females) accumulate DHSM (Figure 4A and B).

**Figure 4:**
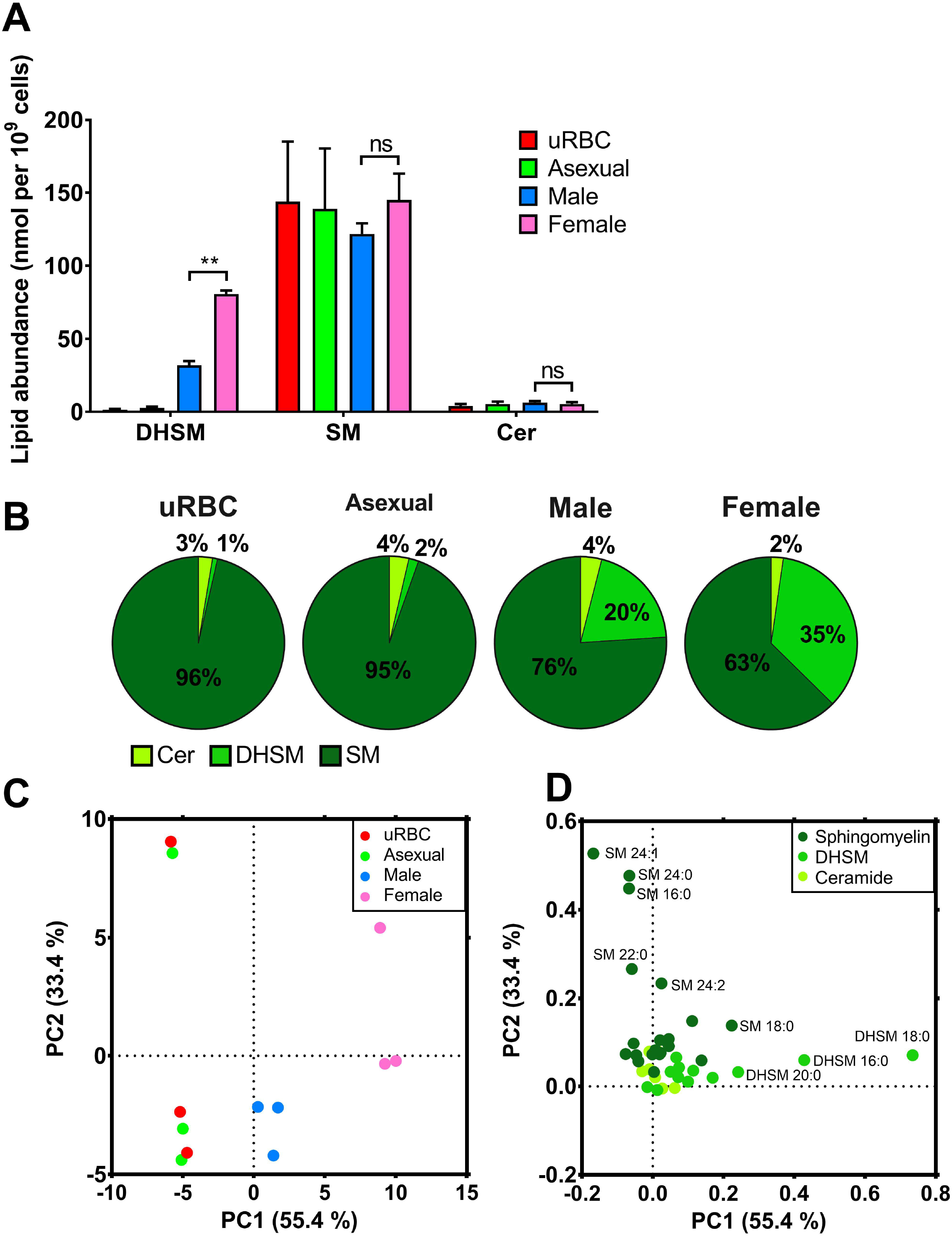
A-B: Mean sphingolipid abundance with standard deviation (A) and mean proportion (B) in uninfected red blood cells (uRBC) and red blood cells infected with asexual stage parasites (Asexual), male gametocytes (Male) or female gametocytes (Female). Results averaged from three independent biological replicates of 10^7^ cells. Significance calculated by two-way ANOVA. **: p<0.01, ns: not significant, p>0.1. C-D: Principal component analysis (C) and loading plot (D) of sphingolipid composition of uninfected red blood cells (uRBC) and red blood cells infected with asexual stage parasites (Asexual), male gametocytes (Male) or female gametocytes (Female). PC1 and 2: first and second principal component; DHSM: dihydrosphingomyelin; SM: sphingomyelin; Cer: ceramide.

Principal component analysis of sphingolipids alone does not distinguish uninfected RBC from asexual parasite-infected RBC. However, male and female gametocyte-infected RBC cluster separately (Figure 4C). DHSM 20:0, 16:0 and 18:0 especially distinguish gametocyte-infected RBC from the other samples (Figure 4D). Overall, accumulation of DHSM marks gametocytogenesis and is even more pronounced in females.

Of the lipid species with more than a two-fold difference between female and male gametocyte-infected RBC, six are significantly more abundant in female gametocyte-infected RBC (Figure 5A). Five of these are saturated DHSM species (between C16-C20 in size), the other is SM 22:2, identifying sphingolipids as the major sex-specific lipid group. On average several neutral lipid species also appear more abundant in female gametocytes, however this difference lacks statistical significance due to variation between biological replicates. No lipid species are significantly more abundant in male gametocyte-infected RBC, although some phospholipid species are more abundant on average. Overall, the sex-specific lipid profile of gametocyte-infected RBC points towards DHSM as a key characteristic of female gametocytes.

**Figure 5:**
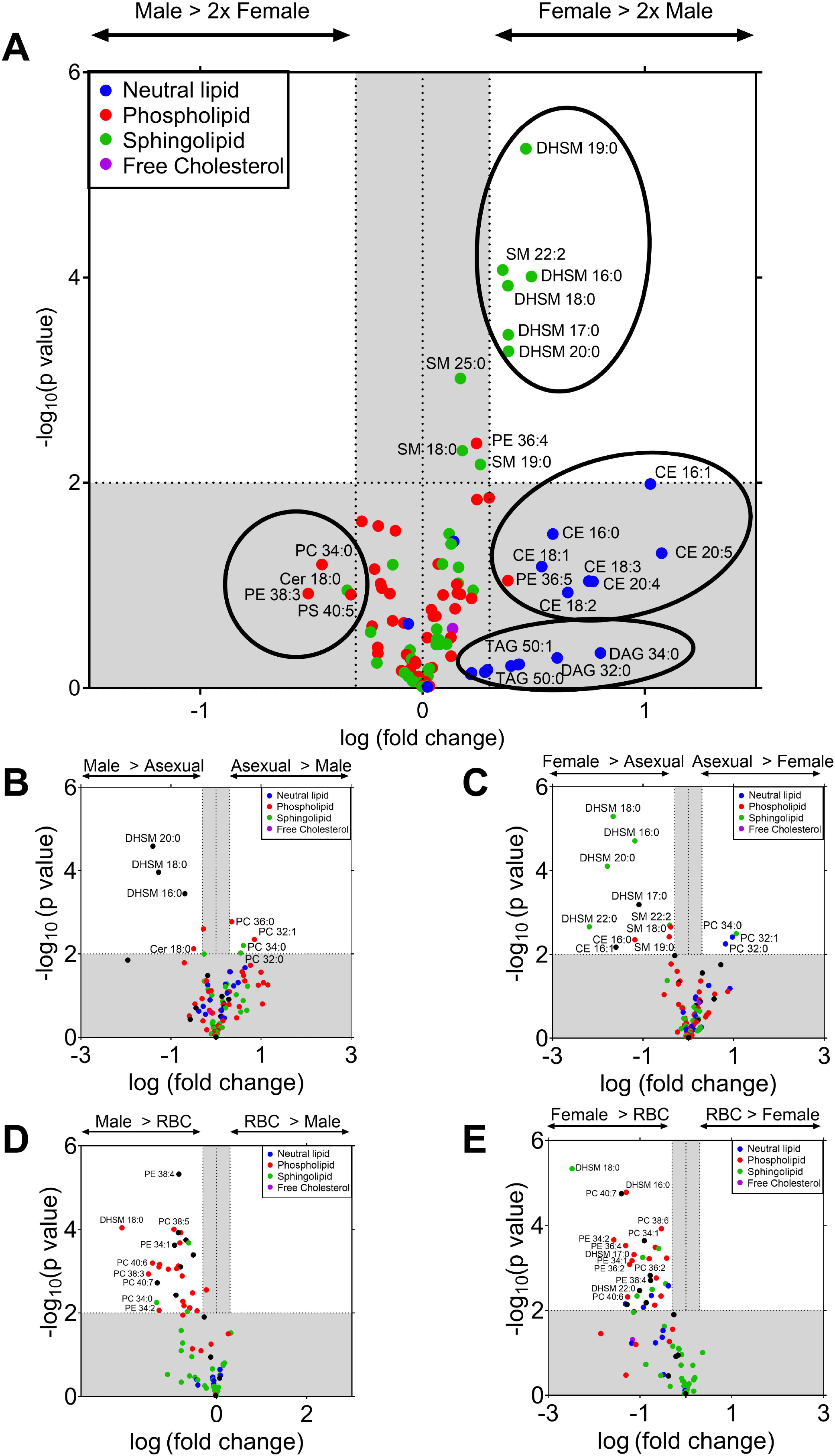
Volcano plot of lipid species in male and female gametocyte-infected red blood cells. A, C and E: Comparison of female gametocyte-infected red blood cell lipids to male gametocyte-infected red blood cells (A), asexual-stage infected red blood cells (C) and uninfected red blood cells (E). B and D: Comparison of male gametocyte-infected red blood cell lipids to asexual-stage infected red blood cells (B) and uninfected red blood cell (RBC) (D). Fold change averaged from three biological repeats of 10^7^ cells each. P values calculated by student t tests. Grey area represents changes that are less than two-fold different between samples and/or have a significance of p> 0.01. Circles in A highlight groups of lipids. DAG: diacylglycerol; DHSM: dihydrosphingomyelin; CE: cholesteryl ester; PC: phosphatidylcholine; PE: phosphatidylethanolamine; PS: phosphatidylserine; SM: sphingomyelin; TAG: triacylglycerol.

Compared to asexual stage parasite-infected RBC, both sexes of gametocyte-infected RBC contain more DHSM 20:0, 18:0 and 16:0 (Figure 5B & C). Asexual parasite-infected RBC on the other hand contain significantly more of some PC species compared to both sexes of gametocyte-infected RBC (PC32:0, PC32:1, PC34:0, PC36:0 [in males only]). CE 16:1 and CE 16:0 are the only lipid species that are more abundant in female gametocyte-infected RBC but not in males when compared to asexual parasite-infected RBC, which might point to specialised functions of these lipid species in female parasites. Several phospholipid and sphingolipid species are significantly more abundant in each sex of gametocyte-infected RBC compared to uninfected RBC (Figure 5D & E). Overall, gametocytes significantly modify the lipid profile of the host RBC during maturation in a manner that distinguishes them from asexual parasites.

### Sphingolipid synthesis is functionally important in both asexual and sexual blood stages

To investigate the functional significance of the sex-specific lipid composition, gametocytes were exposed to three compounds that block the synthesis of the most sex specific lipids (CE and sphingolipids) from day 2 to 6 post commitment (Figure 6). Thereafter the sex-specific viability of gametocytes was measured by flow cytometry. GT11 is an analogue of dihydroceramide that cannot be metabolised by dihydroceramide desaturase in mammalian cells, thereby blocking the enzyme responsible for ceramide synthesis (Bedia et al. 2005). At concentrations greater that 5 μM, GT11 more generally decreased *de novo* sphingolipid synthesis in cultured mammalian cells (Triola et al. 2004). Glibenclamide and Sandoz 58-035 directly inhibit cholesterol esterification by acyl-coenzyme A: cholesterol acyltransferase in mammalian cells (Ross et al. 1984). At 10 *μ* M only the sphingolipid synthesis inhibitor GT11 appears to impact gametocyte viability in a male-specific manner (Figure 6).

**Figure 6:**
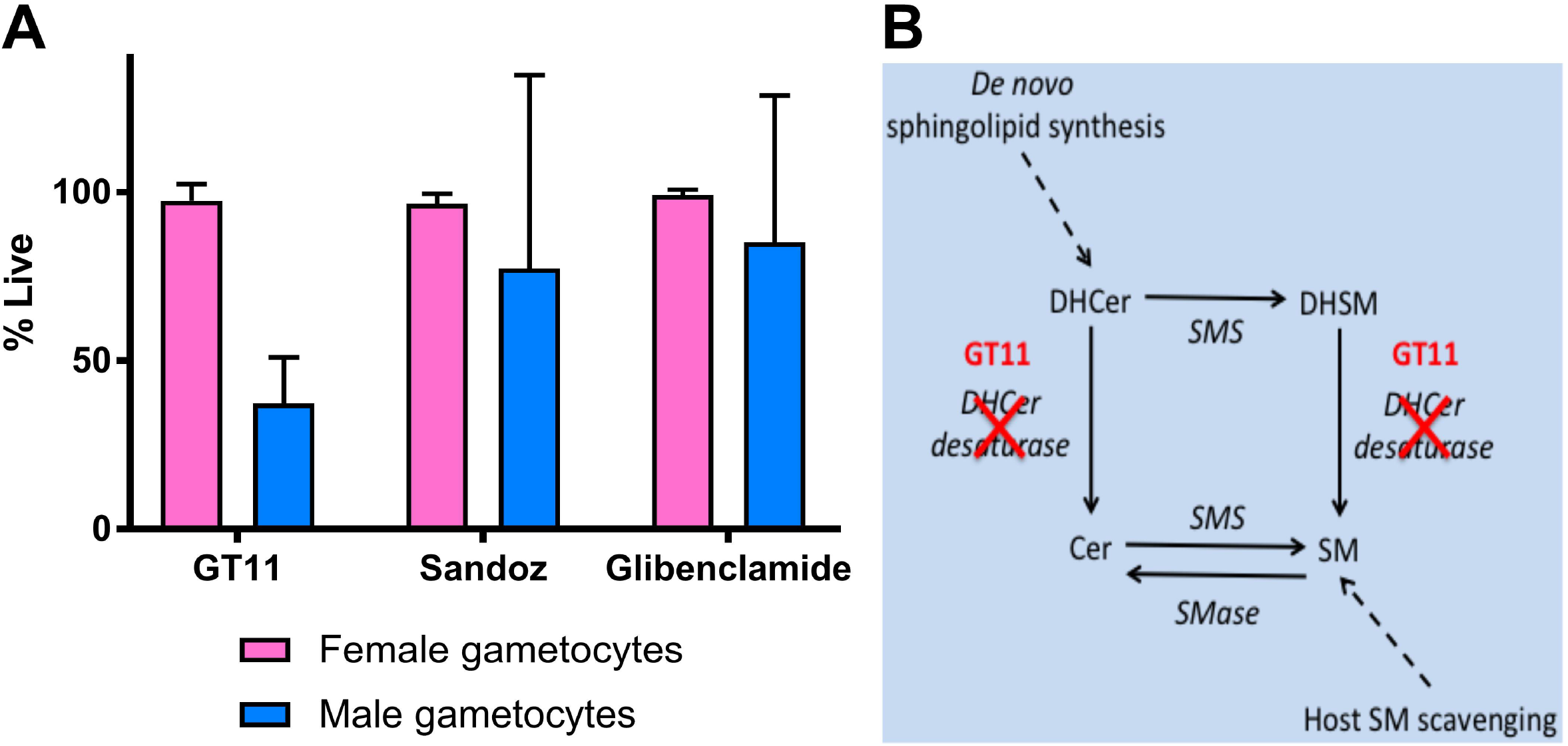
A: Gametocyte viability following chemical inhibition of sphingolipid or CE synthesis. Sphingolipid synthesis was disrupted with 10 μM GT11 and CE synthesis was inhibited with 10 μM Sandoz 58-035 or 10 μM glibenclamide from day 2 to 6 post-commitment. For each gametocyte sex, results are normalised to viability following exposure to the DMSO solvent control and 100 μM artemisinin. Average result and standard deviation presented from three biological replicates, each performed in technical triplicates. B: diagram of GT11 inhibition of sphingolipid synthesis. DHCer: dihydroceramide; DHSM: dihydrosphingomyelin; SMS: sphingomyelin synthase; Cer: ceramide; SM: sphingomyelin.

The apparent sex-specific function of sphingolipid synthesis in gametocytes was further investigated by reverse genetics. DHSM and SM are synthesised by sphingomyelin synthase (SMS) from dihydroceramide and ceramide respectively. *P. falciparum* encodes two such enzymes (SMS1 and SMS2) that are expressed in a parasite stage- and sex-specific manner. SMS activity contributes to the synthesis of SM, a component of detergent-resistant lipid domains in cell membranes, but also catabolises the intracellular signalling molecule ceramide.

Three independent attempts to disrupt both SMS1 and SMS2 genes together did not yield any parasites after transfection. However, disruption of each SMS gene individually produced asexual blood stage parasites that could be analysed further (Supplementary Figures 1 and 2). We first tested the relative fitness of these knock out (KO) parasites. In 28-day co-cultures wild type parasites quickly outgrew each of the KO parasites (Figure 7), suggesting that SMS1 and SMS2 are each required for optimal asexual parasite proliferation. Depleting the culture medium lipids by 75% did not exacerbate the growth defect observed in each KO cell line (Figure 7). In other words, SMS disruption is not readily compensated by an increase in extra-cellular lipid scavenging. Treatment with the sphingolipid synthesis inhibitors GT11 had the same effect on WT and each KO cell line (Figure 8). This suggests that GT11, an analogue of dihydroceramide, does not block SMS in asexual blood stage parasites.

**Figure 7:**
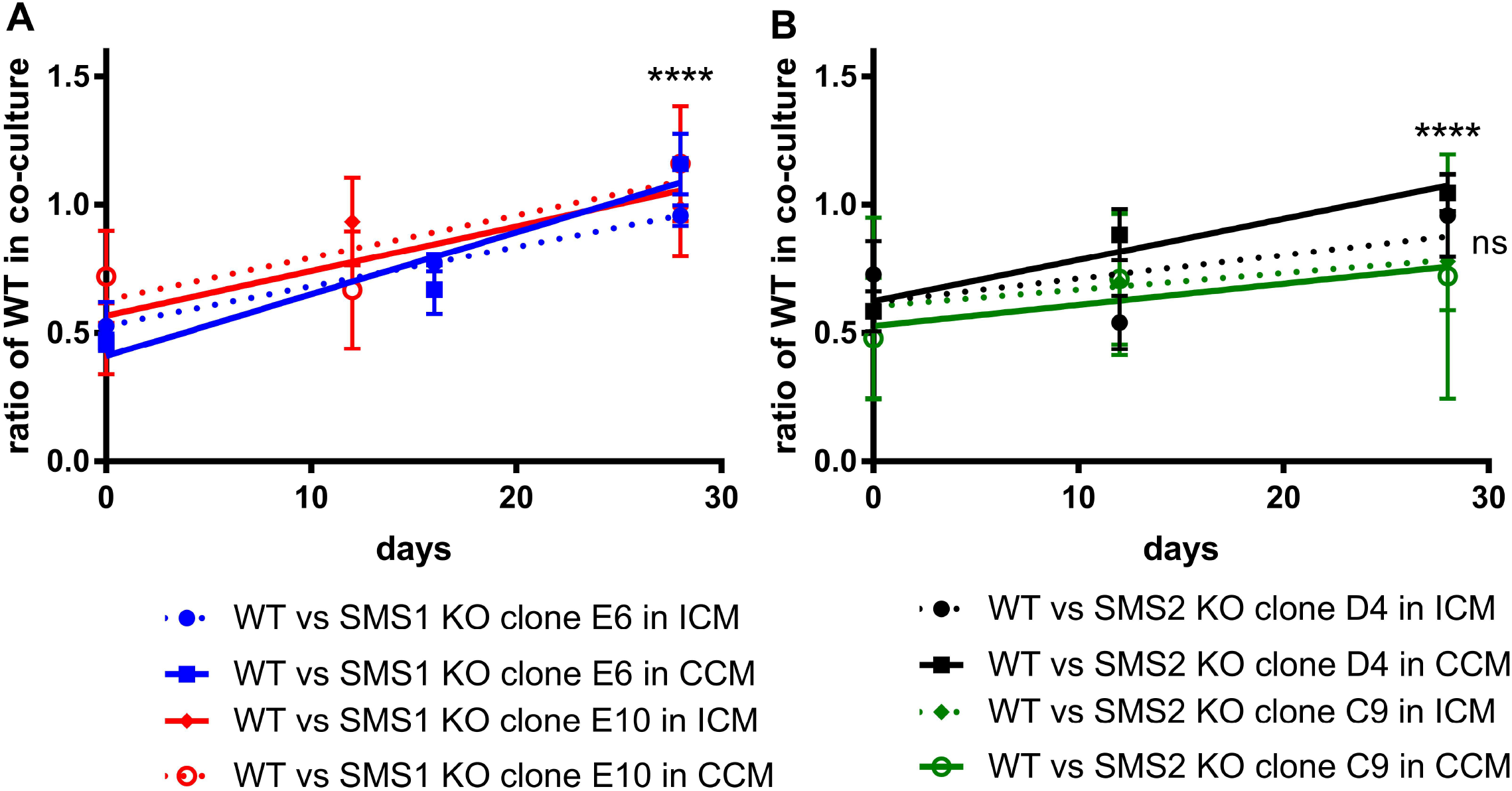
Fitness competition between 3D7 WT and SMS1 KO (A) or SMS2 KO clones (B) in complete (CCM, continuous line) and lipid-depleted culture medium (ICM, dashed line) over 28 days. On day 0 equal amounts of 3D7 WT and SMS1 KO clone E6 (blue, A), SMS1 KO clone E10 (red, A), SMS2 KO clone C9 (green, B) or SMS2 KO clone D4 (black, B) were combined in a co-culture. The proportion of each genotype in the culture was monitored on day 0, 12 and 28 by qPCR of the hDHFR resistance cassette (to detect KO genotype) and the disrupted section of the SMS1 or 2 genes (to detect WT genotype). At some time points technical problems inhibited monitoring the culture so measurements were performed 4 days afterwards instead. Results from 2-4 independent biological repeats are shown as mean and standard deviation. Statistical significance between day 0 and day 28 was calculated by two-way ANOVA. In B results in ICM were not significant (ns), whereas p<0.001 (***) for WT vs SMS1 clone E10 in ICM in A and p<0.0001 (****) for all other results in A and B.

**Figure 8:**
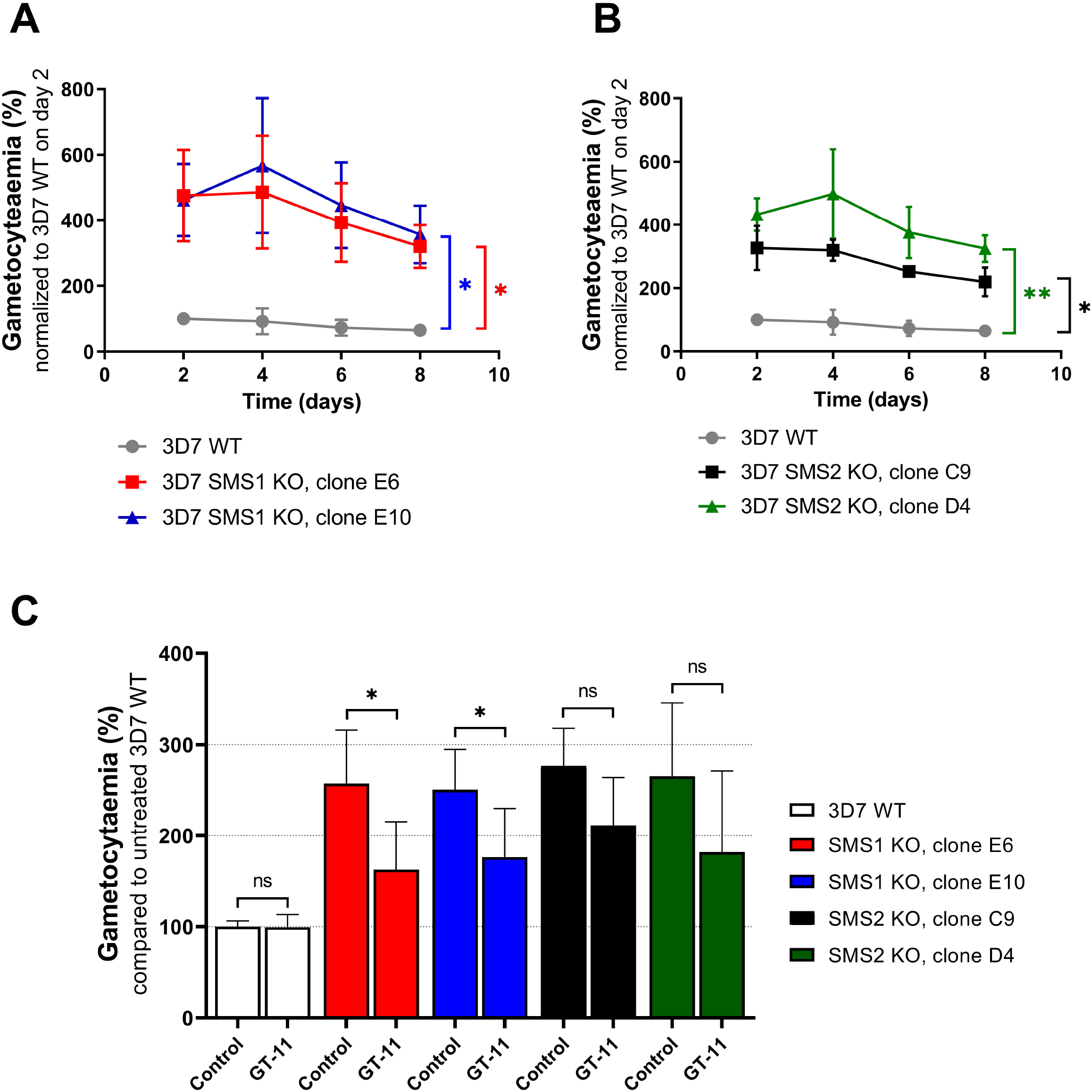
Gametocytaemia in 3D7 WT and SMS1 KO (A) or SMS2 KO clones (B) was monitored every second day from day 2 to day 8 post commitment and compared to WT gametocytaemia on day 2. (C) Effect of 10 μM GT11 on the commitment to gametocytaemia in 3D7 WT, SMS1 KO and SMS2 KO clones. The cultures were treated with GT11 (10 μM) or 0.1% DMSO (control) for 72 hours from day 3 to day 0 pre-commitment. Gametocytaemia was determined on day 8 and compared to gametocytaemia of the untreated WT culture. Gametocytaemia is presented as mean and standard error of mean from three biological replicates, each performed in technical triplicates. Significance calculated by unpaired t-test. ns: not significant, *p*>0.1, *: *p*<0.05, **: *p*<0.01.

**Figure 9:**
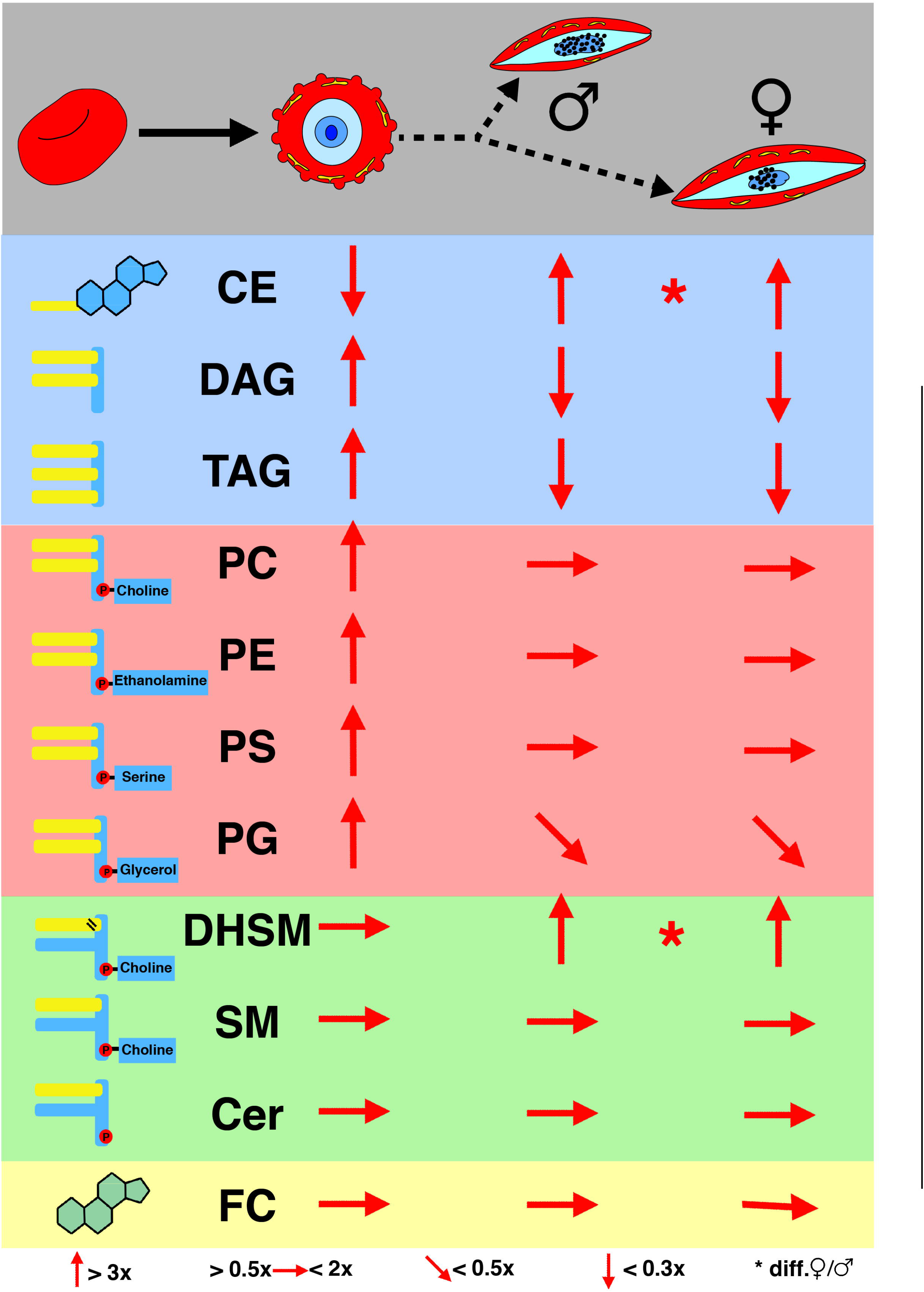
Overview of lipid changes in *P. falciparum* during asexual and sexual blood stages. Grey background: graphic representation of uninfected RBC, RBC infected with trophozoite parasites and male and female infected RBCs. Dashed arrow indicate that there is no direct development from trophozoites to sexual stages and the intermediate stages are not shown. Blue background: neutral lipids; red background: phospholipids; green background: sphingolipids; yellow background: free cholesterol. The left column shows graphic representations of the lipid groups and their abreviation. The red arrows indicate changes relative to the previous life-cycle stage. Upward arrow means >3-fold increase; horizontal arrow means moderate or no changes (changes that are more than 0.5-fold and less than 2-fold); a declining arrow indicates decrease of less than half-fold and a downward arrow signifies a decrease of more than 0.3 of the previous value. The male and female infected RBCs are relative to the trophozoite-infected RBCs. Stars indicate significant changes between male and females. CE: cholesteryl ester; DAG: diacylglycerol; TAG: triacylglycerol; PC: phosphatidylcholine; PE: phosphatidylethanolamine; PS: phosphatidylserine; PG: phosphatidylglycerol; DHSM: dihydrosphingomyelin; SM: sphingomyelin. Cer: ceramide; FC: free cholesterol.

Given that sphingolipid composition is distinctive in gametocytes, the impact of disrupting each SMS was next investigated in sexual blood-stage parasites. The absence of either SMS1 or SMS2 increased gametocyte commitment compared to WT parasites (Figure 8A and B). This may result from asexual blood-stage parasite stress given that gametocyte commitment is a generic parasite stress response (Dixon et al. 2008). When treated with GT11 however, gametocytes lacking a SMS were hypersensitive compared to WT gametocytes (Figure 8C). This suggests that SMS and dihydroceramide desaturase function is additive in gametocytes, especially in conditions where SMS function is limited either through disruption of one of the SMS genes (Figure 8C) or by native reduction of SMS gene expression in males relative to females (Figure 6).

## Discussion

In summary the lipid composition and abundance of lipids in *P. falciparum* gametocytes is sex-specific. Male/female gametocyte lipid dimorphism precedes gametocyte activation and reflects their sex-specific functions.

Female gametocyte-infected RBC stockpile the neutral lipid CE but were resistant to CE synthesis inhibitors Sandoz 58-035 and Glibenclamide. This suggests that acylcoenzyme A: cholesterol acyltransferase-mediated CE synthesis is not essential in mature gametocytes. CE could instead be synthesised by lecithin–cholesterol acyltransferase, or could be scavenged from the host. Alternatively, CE may only be required for later developmental stages. Indeed the lipids in the fertilised zygote are exclusively contributed by the female gametocyte. While the zygote develops in the midgut lumen, neutral lipids are diverted from the blood meal to the eggs of the mosquito *via* lipophorin (Costa et al. 2018). The zygote is therefore vulnerable to lipid depletion in the mosquito midgut lumen, and may depend on maternal CE reserves at this lifecycle stage. In particular, given that mosquitoes are incapable of cholesterol synthesis (Clayton 1964), CE could be a source of cholesterol for the parasite in mosquito stages. *P. falciparum* encodes esterases that could potentially release cholesterol from CE stores (Butler et al. 2020). Neutral lipid staining of the female gametocyte previously highlighted a large neutral lipid body in the cytoplasm (Tran et al. 2014). Potentially the female specific CE store is amassed by the putative lipid transporter gABCG2. Hence, it would be interesting to test the effect of CE synthesis inhibitors on the mosquito stages in future studies.

Male gametocyte-infected RBC do not contain more phospholipids than female gametocyte-infected RBC. Phospholipids are the main component of membranes and are likely required for the rapid cell divisions during male gametocyte activation when eight microgametes are formed within the male gametocyte. While additional plasma membranes may not be required in females, female gametocytes contain an elaborate mitochondrial network bound by a double membrane rich in phospholipids, as well as abundant membrane bound osmiophilic bodies (Langreth et al. 1978; Jensen 1979; Sinden 1982; Ponnudurai et al. 1986). Although the total phospholipid content of male and female gametocyte-infected RBC is similar, the allocation of phospholipids between organelles may be sex-specific.

For example, PG serves as the precursor for cardiolipin found exclusively in the mitochondria (Gebert et al. 2009; de Kroon et al. 1999; Daum 1985). Consistent with this, PG is absent in uninfected RBC, which lack mitochondria. Surprisingly however, there is no difference in PG between male and female gametocytes despite female gametocytes relying more on their mitochondria than males. Indeed mitochondrial proteins have been shown to be more abundant in female than in male gametocytes (Miao et al., 2017). Female gametes in the mosquito have an increased energy demand (probably in anticipation of the post-fertilisation stage (MacRae et al. 2013), whereas male gametocytes lose their mitochondria completely during the development into microgametes (Okamoto et al. 2009). The presence of comparable amounts of PG both in male and female iRBC argues for a similar number or size of mitochondria at least at the analysed stage of development (stage IV gametocytes). However, less PG is detected in the RBC infected with sexual stages compared to the RBC infected with asexual stages despite the upregulation of tricarboxylic acid cycle function in gametocytes (MacRae 2013). This cautions against the simple correlation between PG and mitochondrial function. Nonetheless, the specificity and accumulation of PG in the parasite could be targeted by antimalarial compounds effective against both asexual and sexual blood stage parasites.

Phospholipid composition distinguished both male and female gametocyte-infected RBC from asexual parasite-infected RBC and uninfected RBC. Less phospholipids were present in gametocyte-infected RBC compared to asexual parasite-infected RBC consistent with asexual replication requiring phospholipid-rich membrane biogenesis (e.g. for the plasma membrane of a growing parasites, for merozoite formation and for RBC modifications such as Maurer’s clefts). The increase in PS between trophozoites and gametocytes reported by Gulati et al. (2015) was not observed in this study, perhaps due to technical differences in sample preparation.

In asexual blood stages, reduction of sphingolipid metabolism by disruption of either SMS1 or SMS2 was not lethal, but incurred a growth defect. Disruption of both SMS genes on the other hand appears to be lethal for asexual blood stage parasites, despite SMS2 being expressed at very low levels in this lifecycle stage. This suggests that *de novo* sphingomyelin synthesis is essential in asexual blood stages and indicates that each SMS is – to a certain degree - functionally redundant. Surprisingly this defect was not exacerbated by depleting the culture medium lipids by 75%, or by inhibiting dihydroceramide desaturase with GT11. This supports the hypothesis that SMS activity regulates intracellular ceramide concentration rather than sphingomyelin itself being essential for asexual blood stage proliferation. Indeed, ceramide might otherwise build up to cytotoxic levels due to sphingomyelinase activity, which has previously been described in asexual stages (Hanada et al. 2002). Inhibition of ceramide production by sphingomyelinase with GW4869 is also lethal for asexual *P. falciparum* (Gulati et al. 2015), further illustrating the importance of regulating ceramide abundance.

The most distinctive feature of gametocyte-infected RBC, particularly female gametocyte-infected RBC, is the presence of DHSM species. This is consistent with other studies that suggest *de novo* sphingolipid synthesis, rather than host SM hydrolysis by neutral sphingomyelinase, is active in gametocytes (Gulati et al. 2015; Tran et al. 2016). Here we sought to test whether DHSM accumulation in gametocytes results from decreased catabolism by dihydroceramide desaturase and/or from increased synthesis by SMS. Inhibition of dihydroceramide desaturase with 10 μM GT11 selectively killed male gametocytes (Figure 6) suggesting that dihydroceramide desaturase activity is sex-specific, which may contribute to the difference in DHSM abundance. The male-specific effect of GT11 was not observed in combined male and female gametocytes (Figure 8), presumably due to the gametocyte population being female biased. However, gametocytes that lack an SMS were hypersensitive to GT11 (Figure 8). In other words, reduced SMS activity (due to endogenous differential SMS gene expression between male and female gametocytes or through SMS gene disruption) appears to make parasites more susceptible to dihydroceramide desaturase inhibition by GT11. Given that dihydroceramide desaturase and SMS share dihydroceramide as a substrate, hypersensitivity to GT11 in gametocytes with reduced SMS activity may result from an accumulation of dihydroceramide. Conversely, accumulation of DHSM in gametocytes may be a means of reducing intracellular dihydroceramide levels.

Similarly, the observed increased gametocyte commitment in SMS KO cell lines could be a stress response to elevated dihydroceramide concentrations. Indeed it has recently been observed that dihydroceramides are biologically active lipids involved in diverse mammalian cell functions (Siddique et al. 2015). Lipids have previously been implicated in inducing gametocytogenesis in *P. falciparum* (Brancucci et al. 2017). In particular, depletion of serum lysophosphatidylcholine, a precursor of PC synthesis, is known to trigger gametocytogenesis. We observed no significant difference in lysophosphatidylcholine abundance between male and female gametocytes (supplementary figure S3). PC is a substrate of SMS-mediated SM and DHSM synthesis. As such, reducing parasite SMS activity may have mimicked the downstream effect of PC depletion and triggered gametocytogenesis.

Overall, this study has shown that lipid metabolism is not only variable between parasite lifecycle stages, but is also sex-specific. These differences have implications for the effectiveness of drugs targeting metabolic pathways, and should be considered in the design of future transmission blocking antimalarial treatments. In terms of the fundamental biology of the parasite, the lipid profile of gametocytes is a testament to the parasite’s ability to thrive in two radically different host environments. The lipidome presented here provides a reference for unravelling the complex sex-specific function of lipids in the *P. falciparum*.

## Acknowledgements

FACS was performed with assistance from Dr. Harpeet Vohra and Mr. Michael Devoy. We are grateful to the Australian Red Cross for providing human red blood cells and serum. Funding was provided by the Australian Research Council (DP180103212) and the National Health and Medical Research Council of Australia (APP1182369). M.C.R. is supported by the Australian Government Research Training Program Scholarship and The Australian National University.

## Author contributions

M.C.R. and A.G.M. conceived and designed the study; M.C.R., P.T., S.B. and D.C. performed the experiments; M.C.R., S.B., D.C., T.M. and A.G.M. analysed the data; and M.C.R. and A.G.M. wrote the paper. All authors contributed to the editing of the paper.

## Submission of data set to database

The mass spectrometry raw data of the lipidomics experiments have been deposited in the MetaboLights database (http://www.ebi.ac.uk/metabolights).

