## Supplemental Material for "Analysis of sex-specific lipid metabolism in *P. falciparum* gametocytes points to importance of sphingomyelin for gametocytogenesis"

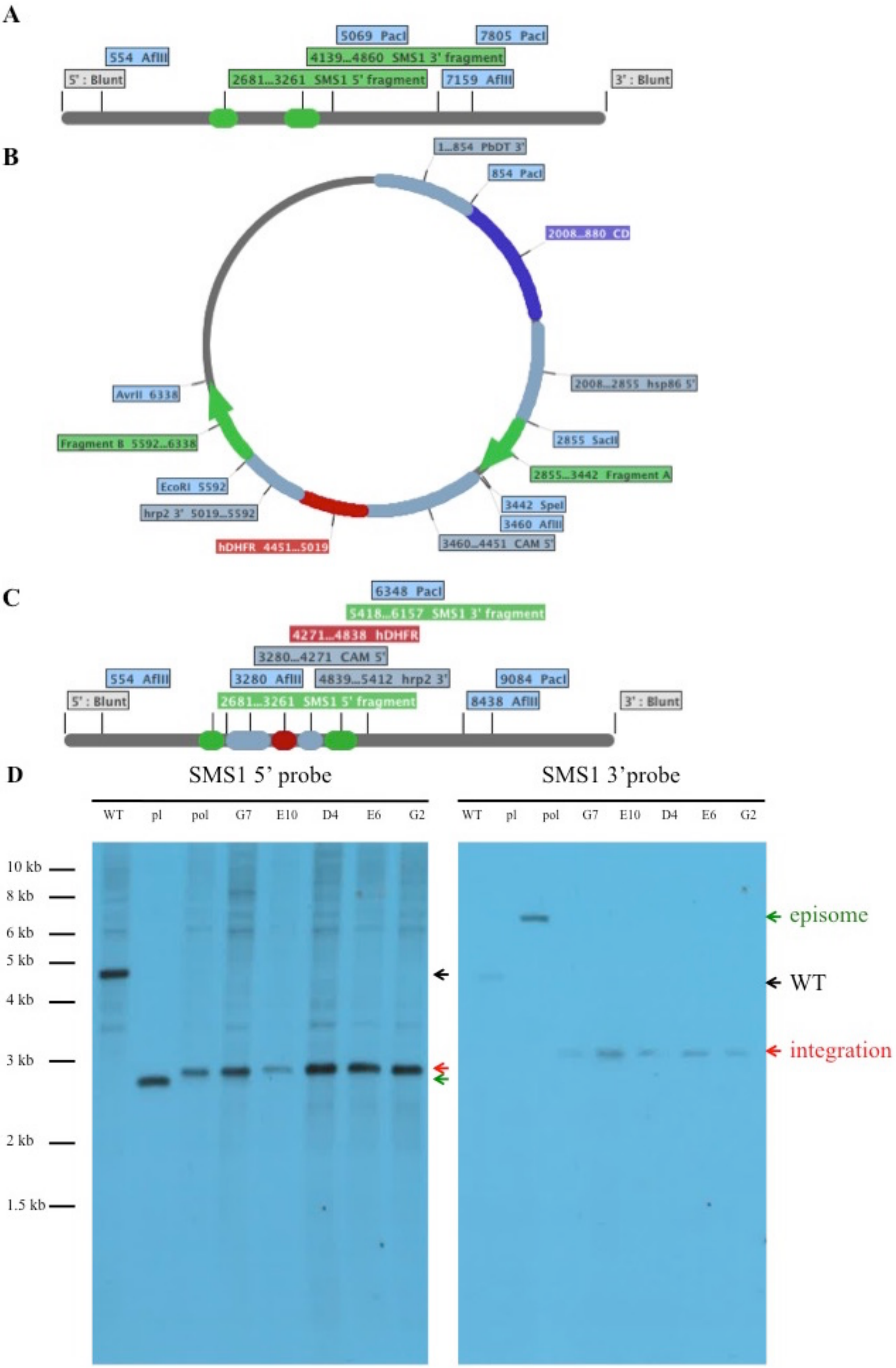
A
5069 PacI
7805 PacI
554 AflII
4139...4860 SMS1 3' fragment
5' : Blunt
2681...3261 SMS1 5' fragment
7159 AflII
3' : Blunt
B
1...854 PbDT 3'
854 PacI
2008...880 CD
2008...2855 hsp86 5'
2855 SacII
2855...3442 Fragment A
3442 SpeI
3460 AflII
3460...4451 CAM 5'
hDHFR 4451...5019
hrp2 3' 5019...5592
EcoRI 5592
Fragment B 5592...6338
AvrII 6338
C
6348 PacI
5418...6157 SMS1 3' fragment
4271...4838 hDHFR
3280...4271 CAM 5'
554 AflII
3280 AflII
4839...5412 hrp2 3'
9084 PacI
5' : Blunt
2681...3261 SMS1 5' fragment
8438 AflII
3' : Blunt
D
SMS1 5' probe
SMS1 3'probe
WT pl pol G7 E10 D4 E6 G2
WT pl pol G7 E10 D4 E6 G2
10 kb
8 kb
6 kb
5 kb
4 kb
3 kb
2 kb
1.5 kb
episome
WT
integration

**Supplementary Figure 1:** Disruption of SMS1 by homologous recombination. A-C) Schematic of WT SMS1 locus with surrounding fragment of genomic DNA (A), pCC-1/SMS1 plasmid (B) and disrupted SMS1 locus with surrounding fragment of genomic DNA (C). SMS1 gene is disrupted by double homologous recombination at the 5’ and 3’ extremities of the gene in green (SMS1 5’ fragment between SacII and SpeI restriction sites and SMS1 3’ fragment between EcroRI and AvrII restriction sites), which inserts positive selection marker human dihydrofolate reductase (hDHFR, red) under the control of the calmodulin promoter (CAM 5’) and histidine rich protein 2 3’ untranslated region (hrp2 3’). The plasmid also contains negative selection maker cytosine deaminase (CD, purple) under the control of heat shock protein 86 promoter (hsp86 5’) and *P. berghei* dihydrofolate reductase 3’ untranslated region (PbDT 3’) that are lost by double recombination. AflII and PacI restriction digest sites used for Southern blot are also indicated. D) Southern blot showing disruption of SMS1 locus. DNA from 3D7 wild type (WT), SMS1 KO plasmid (pl), polyclonal 3D7 transfected with SMS1 KO plasmid (pol) or clonal 3D7 transfected with SMS1 KO plasmid (G7, E10, D4, E6, G2) digested with AflII and PacI restriction enzymes and probed with 5’ homologous flank probe (SMS1 5’ probe, left) or 3’ homologous flank probe (SMS1 3’ probe, right). Expected SMS1 5’ probe band lengths: WT locus: 4,515 bp (black arrow), episomal locus: 2,606 bp (green arrow), integrated locus: 2,726 bp (red arrow). Expected SMS1 3’ probe band lengths: WT locus: 4,515 bp (black arrow), episomal locus: 6,243 bp (green arrow), integrated locus: 3,068 bp (red arrow).

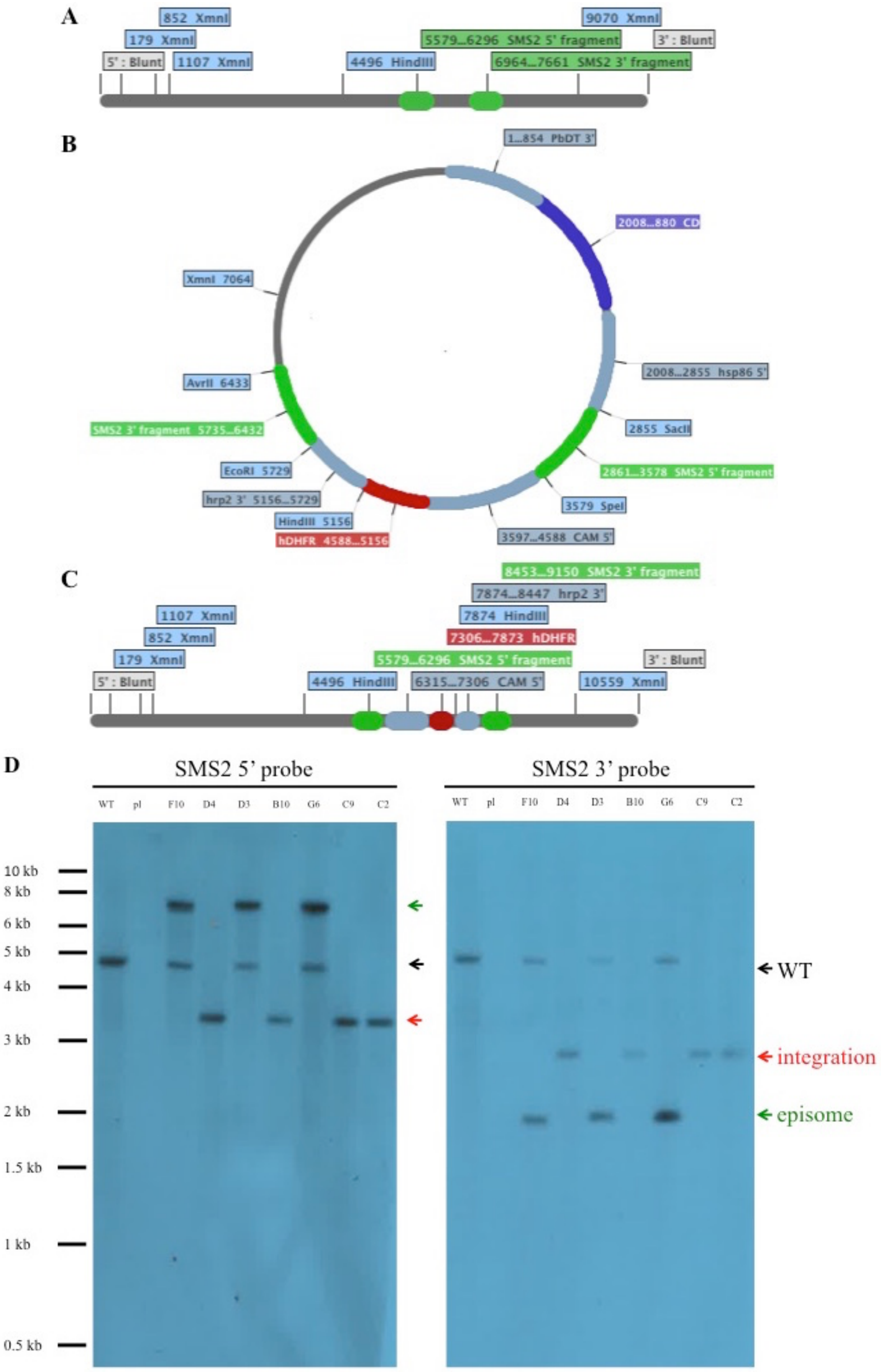

A
852 XmnI
179 XmnI
5' : Blunt
1107 XmnI
4496 HindIII
5579...6296 SMS2 5' fragment
6964...7661 SMS2 3' fragment
9070 XmnI
3' : Blunt
B
1...854 PbDT 3'
2008...880 CD
2008...2855 hsp86 5'
2855 SacII
2861...3578 SMS2 5' fragment
3579 SpeI
3597...4588 CAM 5'
hDHFR 4588...5156
HindIII 5156
hrp2 3' 5156...5729
EcoRI 5729
SMS2 3' fragment 5735...6432
AvrII 6433
XmnI 7064
C
8453...9150 SMS2 3' fragment
7874...8447 hrp2 3'
7874 HindIII
7306...7873 hDHFR
5579...6296 SMS2 5' fragment
6315...7306 CAM 5'
4496 HindIII
1107 XmnI
852 XmnI
179 XmnI
5' : Blunt
3' : Blunt
10559 XmnI
D
SMS2 5' probe
SMS2 3' probe
WT
pl
F10
D4
D3
B10
G6
C9
C2
10 kb
8 kb
6 kb
5 kb
4 kb
3 kb
2 kb
1.5 kb
1 kb
0.5 kb
WT
integration
episome

**Supplementary Figure 2:** Disruption of SMS2 by homologous recombination. A-C) Schematic of WT SMS2 locus with surrounding fragment of genomic DNA (A), pCC-1/SMS2 plasmid (B) and disrupted SMS2 locus with surrounding fragment of genomic DNA (C). SMS2 gene is disrupted by double homologous recombination at the 5' and 3' extremities of the gene in green (SMS2 5' fragment between SacII and SpeI restriction sites and SMS2 3' fragment between EcroRI and AvrII restriction sites), which inserts positive selection marker human dihydrofolate reductase (hDHFR, red) under the control of the calmodulin promoter (CAM 5') and histidine rich protein 2 3' untranslated region (hrp2 3'). The plasmid also contains negative selection maker cytosine deaminase (CD, purple) under the control of heat shock protein 86 promoter (hsp86 5') and *P. berghei* dihydrofolate reductase 3' untranslated region (PbDT 3') that are lost by double recombination. XmnI and HindIII restriction digest sites used for Southern blot are also indicated. D) Southern blot showing disruption of SMS2 locus. DNA from 3D7 wild type (WT), SMS2 KO plasmid (pl), or clonal 3D7 transfected with SMS2 KO plasmid (G7, E10, D4, E6 and G2) digested with XmnI and Hind III restriction enzymes probed with 5' homologous flank probe (SMS2 5' probe, left) or 3' homologous flank probe (SMS2 3' probe, right). Expected SMS2 5' probe band lengths: WT locus: 4,574 bp (black arrow), episomal locus: 7,036 bp (green arrow), integrated locus: 3,378 bp (red arrow). Expected SMS2 5' probe band lengths: WT locus: 4,574 bp (black arrow), episomal locus: 1,908 bp (green arrow), integrated locus: 2,685 bp (red arrow).

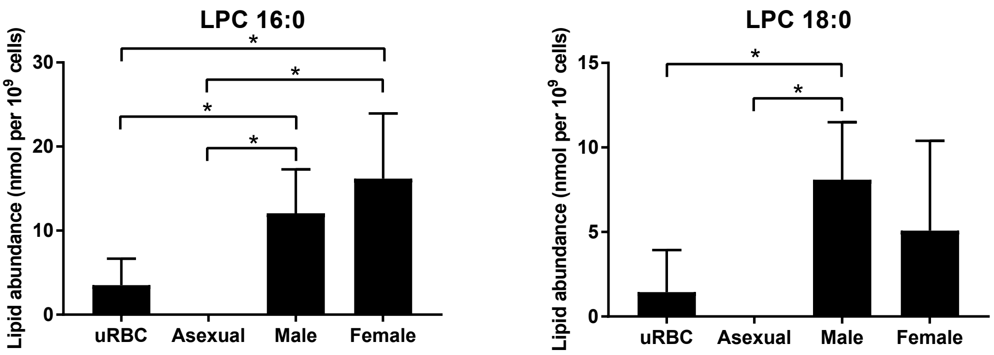


**Supplementary figure 3:** Abundance of lysophosphatidylcholine (LPC) 16:0 (left) and LPC 18:0 (right) in uninfected red blood cells (uRBC) and red blood cells infected with asexual stage parasites (Asexual), male gametocytes (Male) or female gametocytes (Female) with results from student t tests (*: $p<0.1$; nothing indicated where $p>0.1$).

**Supplementary Table 1:** Primers for construct preparation. Lowercase letters in the primer sequences correspond to restriction enzyme sites while uppercase letters refer to the gene sequence.

| Primer code | Primer description | | Primer sequence |
|---|---|---|---|
| al398 | SMS1 5’ | forward | atcccgcggCACACATTTGTACCTCTC |
| al399 | | reverse | gatactagtATCTGAGAAATTGGAACGC |
| al400 | SMS1 3’ | forward | atcgaattcGCTGCAAGAAGATATGC |
| al401 | | reverse | gatcctaggAAAAAGAGTTTGTAGGTG |
| al406 | SMS2 5’ | forward | atcccgcggGTTTAATACACGTGAG |
| al407 | | reverse | gatactagtCGATCACTTAATGGTTGCG |
| al408 | SMS2 3’ | forward | atcgaattcTATACCTTAGATTATGCC |
| al409 | | reverse | gatcctaggAAAAAACGACATTTAGGG |

**Supplementary Table 2:** Fitness competition primers to quantify wild type (SMS1 or SMS2 “cut out”) and knock out (hDHFR) parasite genotype abundance relative to total parasite abundance (reference).

| Target | Gene ID | Primers (forward/reverse) |
|---|---|---|
| SMS1 “cut out” | PF3D7_0625000 | ACTGTTGATGTGTTAATGGGATATG<br>ATCTTCTTGCAGCTACATCTACTA |
| SMS2 “cut out” | PF3D7_0625100 | TTCATGCAAAGCCATTCTTTCT<br>TGGCATAATCTAAGGTATAATTCAATCC |
| hDHFR | | GAACTCAAGGAACCTCCACAA<br>ACAGAACTGCCACCAACTATC |
| Reference | PF3D7_0317300 | AATAGTCGAAGCGGGAAGTG<br>CGAATTGGATTCTCCCAAATAACC |

**Supplementary Table 3:** Primers for Southern blot probe synthesis

| Primer code | Primer description | | Primer sequence |
|---|---|---|---|
| al 514 | SMS1 5’ | forward | CACACATTTGTACCTCTCTTA |
| al515 | | reverse | CCTTATGGTTGTAATGTTTGTC |
| al516 | SMS1 3’ | forward | TCACAGACAGATTCCAAGTGTT |
| al517 | | reverse | GGTACCATTTCCAGCGTATGA |
| al518 | SMS2 5’ | forward | GTAAATTAAAGACATGCACGTGAGA |
| al519 | | reverse | CGCTATCTGTTACTGACTCTTCAT |
| al520 | SMS2 3’ | forward | ACCCATAACAGAGGATAA |
| al521 | | reverse | ACGACATTTAGGGATTAAA |

**Supplementary Table 4:** list of detected lipid species other than free cholesterol in *P. falciparum*. CE: cholesteryl ester; DAG: diacylglycerol; TAG: triacylglycerol; LPC: lyso-phosphatidylcholine; PC: phosphatidylcholine; PE: phosphatidylethanolamine; PG: phosphatidylglycerol; PS: phosphatidylserine; Cer: ceramide; DHSM: dihydrosphingomyelin; SM: sphingomyelin.

| **Neutral Lipids** | | |
|---|---|---|
| CE 16:0 | DAG (34:0) 16:0_18:0 | TAG 52:1 |
| CE 16:1 | DAG (34:1) 16:0_18:1 | TAG 52:2 |
| CE 18:1 | DAG (36:1) 18:0_18:1 | TAG 52:3 |
| CE 18:2 | DAG (36:2) 18:1_18:1 | TAG 54:2 |
| CE 18:3 | TAG 48:0 | TAG 54:3 |
| CE 20:4 | TAG 50:0 | TAG 54:6 |
| CE 20:5 | TAG 50:1 | TAG 56:6 |
| DAG (32:0) 16:0_16:0 | TAG 50:2 | |

| Phospholipids | | |
|---|---|---|
| LPC 16:0 | PC 38:7 | PE 38:4 |
| LPC 18:0 | PC 40:4 | PE 38:5 |
| PC 32:0 | PC 40:5 | PE 38:6 |
| PC 32:1 | PC 40:6 | PE 40:5 |
| PC 34:0 | PC 40:7 | PE 40:6 |
| PC 34:1 | PC O-32:0 | PG (34:1) 16:0_18:1 |
| PC 34:2 | PC O-34:1 | PG (34:2) 16:0_18:2 |
| PC 34:3 | PC O-36:2 | PG (36:1) 18:0_18:1 |
| PC 36:0 | PC O-38:4 | PG (36:2) 18:0_18:2 |
| PC 36:1 | PC O-38:5 | PG (36:2) 18:1_18:1 |
| PC 36:2 | PE 34:1 | PG (36:3) 18:1_18:2 |
| PC 36:3 | PE 34:2 | PS 34:1 |
| PC 36:4 | PE 36:1 | PS 36:1 |
| PC 36:5 | PE 36:2 | PS 36:2 |
| PC 38:3 | PE 36:3 | PS 38:4 |
| PC 38:4 | PE 36:4 | PS 38:5 |
| PC 38:5 | PE 36:5 | PS 40:5 |
| PC 38:6 | PE 38:3 | PS 40:6 |

| Sphingolipids | | |
|---|---|---|
| Cer 16:0 | DHSM 25:0 | SM 22:2 |
| Cer 18:0 | SM 14:0 | SM 23:0 |
| Cer 19:0 | SM 15:0 | SM 23:1 |
| Cer 22:0 | SM 16:0 | SM 24:0 |
| Cer 24:0 | SM 16:1 | SM 24:1 |
| Cer 24:1 | SM 17:0 | SM 24:2 |
| Cer 24:2 | SM 18:0 | SM 24:3 |
| DHSM 16:0 | SM 18:1 | SM 25:0 |
| DHSM 17:0 | SM 19:0 | SM 25:1 |
| DHSM 18:0 | SM 20:0 | SM 26:0 |
| DHSM 19:0 | SM 20:1 | SM 26:1 |
| DHSM 20:0 | SM 21:0 | SM 26:2 |
| DHSM 22:0 | SM 22:0 | |
| DHSM 24:0 | SM 22:1 | |
